# Bakta: Rapid & standardized annotation of bacterial genomes via alignment-free sequence identification

**DOI:** 10.1101/2021.09.02.458689

**Authors:** Oliver Schwengers, Lukas Jelonek, Marius Dieckmann, Sebastian Beyvers, Jochen Blom, Alexander Goesmann

**Affiliations:** Bioinformatics and Systems Biology, Justus Liebig University Giessen, Giessen, 35392, Germany

**Keywords:** genome annotation, bacteria, plasmids, whole-genome sequencing

## Abstract

Command line annotation software tools have continuously gained popularity compared to centralized online services due to the worldwide increase of sequenced bacterial genomes. However, results of existing command line software pipelines heavily depend on taxon specific databases or sufficiently well annotated reference genomes. Here, we introduce Bakta, a new command line software tool for the robust, taxon-independent, thorough and nonetheless fast annotation of bacterial genomes. Bakta conducts a comprehensive annotation workflow including the detection of small proteins taking into account replicon metadata. The annotation of coding sequences is accelerated via an alignment-free sequence identification approach that in addition facilitates the precise assignment of public database cross references. Annotation results are exported in GFF3 and INSDC-compliant flat files as well as comprehensive JSON files facilitating automated downstream analysis. We compared Bakta to other rapid contemporary command line annotation software tools in both targeted and taxonomically broad benchmarks including isolates and metagenomic-assembled genomes. We demonstrated that Bakta outperforms other tools in terms of functional annotations, the assignment of functional categories and database cross-references whilst providing comparable wall clock runtimes. Bakta is implemented in Python 3 and runs on MacOS and Linux systems. It is freely available under a GPLv3 license at https://github.com/oschwengers/bakta. An accompanying web version is available at https://bakta.computational.bio.

## Introduction

Regional and functional annotations have become a routine task in the analysis of bacterial whole-genome sequencing data. A thorough genome annotation is crucial to form a stable basis for many downstream analyses as both accuracy and comprehensiveness of the annotation have strong impacts on the outcome of related studies. Hence, various online services evolved to streamline the different steps that are involved in this task [1–4]. However, these services have become unsuitable for the timely annotation of high-throughput data which is needed to keep pace with the ever increasing speed at which bacterial genomes are sequenced today [5]. To meet these growing demands, annotations are required to be conducted either locally on standard consumer hardware or within high-performance or cloud computing infrastructures. Therefore, several command line software tools for the rapid annotation of bacterial genomes have recently been developed, *e.g*. Prokka [6] and DFAST [7].

These tools, however, trade annotation database sizes and workflow standardizations for runtime performance and flexibility regarding user-provided annotation data, respectively. In particular, requirements for taxon-specific databases are drawbacks for automated high-throughput annotations in situations where no or only limited taxonomic knowledge is available *a priori*, for instance as part of larger analysis pipelines [8–11]. Likewise, requirements for annotated reference genomes present an obstacle for the annotation of species that are underrepresented in public databases or for which no annotated reference genomes are available, *e.g*. metagenome-assembled genomes (MAGs). Depending on taxonomic groups [12], these are important issues often involved in low rates of functionally described and annotated genes. Furthermore, existing rapid offline annotation software tools leave room for improvements regarding the following issues: (*i*) despite the discovery of previously overlooked conserved short open reading frames (sORFs) two decades ago [13], they neither predict nor detect coding sequences (CDSs) of nowadays well-known small proteins shorter than 29 amino acids, due to technical gene length cutoffs implemented within underlying gene prediction tools to reduce the number of false *de novo* predictions [14,15]; (*ii*) they do not identify known protein sequences stored in public databases like RefSeq [16] and UniRef100 [17] and thus cannot assign database cross references (dbxrefs), *i.e*. stable public database identifiers facilitating the interconnection with further and more detailed databases; (*iii*) they do not take into account additional sequence information, *i.e*. completeness and topology, for the structural annotation of CDSs spanning artificial sequence edges.

Addressing these issues, here we introduce Bakta, a new command line tool for the automated and standardized annotation of bacterial genomes aiming at a well-balanced tradeoff between runtime performance and comprehensive annotations. It implements a comprehensive annotation workflow for coding and non-coding genes complemented by the prediction of CRISPR arrays, gaps, oriC and oriT features. In contrast to other lightweight annotation pipelines, Bakta is able to detect and annotate small proteins by a custom extraction and filter workflow for sORFs. The CDSs annotation workflow is accelerated by a hash-based alignment-free protein sequence identification approach considerably reducing the number of required computationally expensive sequence alignments. This new approach furthermore facilitates the annotation of CDSs with cross references to public databases via stable identifiers. We envision Bakta also as a suitable software tool for integration into larger pipelines. To streamline this process, results and supplementary information are additionally provided as comprehensive and well-structured JSON files.

## Design and implementation

### Annotation workflow

Bakta implements a comprehensive workflow capable of utilizing sequence metadata in addition to the genome assembly. It annotates coding and non-coding genes, CRISPR arrays, gaps, oriC and oriT features (Fig. 1) that are rigorously filtered by annotation information and overlaps. Final results are exported in human and machine readable formats as well as standard bioinformatics file formats. The following sections provide a detailed description of all Bakta annotation workflow steps.

**Figure 1:**
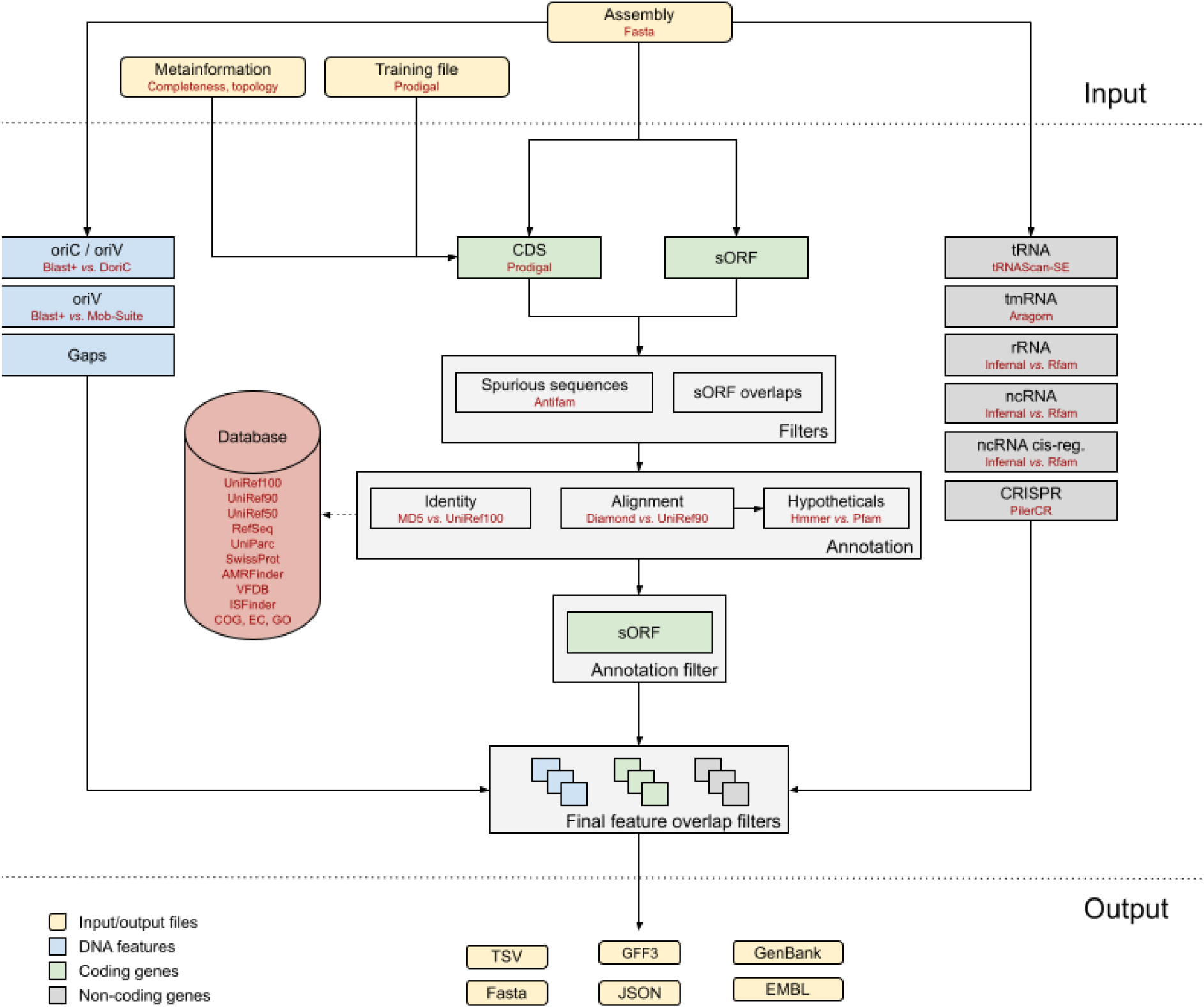
Overview of the Bakta annotation workflow

Bakta accepts assembled genome sequences in optionally zipped Fasta format. To improve the structural annotation of CDSs within finished genomes or complete replicons of draft assemblies, sequence metadata, *e.g*. completeness and topology, can optionally be provided as tab-separated values (TSV). To improve the prediction of CDSs in draft assemblies, precomputed Prodigal [15] training files can be provided as well if available.

Transfer-RNA and transfer-messenger-RNA genes are predicted and annotated by tRNAscan-SE [18] and Aragorn [19], respectively. Ribosomal genes and non-coding RNAs are predicted and annotated by Infernal [20] using Rfam [21] covariance models. It is worth noting that non-coding RNA genes and non-coding RNA cis-regulatory elements are predicted and annotated as distinct feature types, allowing for distinct annotations of regulatory region subtypes and adjusted feature overlap filters. CRISPR arrays are predicted by Piler-CR [22]. Origins of replication and origins of transfer are detected by BLAST+ [23] against sequences from DoriC [24] and MOB-suite [25], respectively.

Coding sequences are predicted by Prodigal taking into account optionally provided metadata on sequence completeness and topology, enabling the prediction of CDSs spanning artificial replicon edges. Therefore, predicted pairs of partial CDSs on complete replicons that run off the 5’ and 3’ edges on the same strand are merged by Bakta. sORFs of small proteins shorter than 30 amino acids are extracted with BioPython [26]. Publicly known spurious CDSs and sORFs are filtered out using HMMER [27] and AntiFam [28] hidden markov models (HMM) of false positive sequences, e.g. shadow open reading frames along tRNA genes. To accelerate the annotation process, publicly known unique protein sequences (UPSs) of CDSs and sORFs are identified via an alignment-free hash-based approach, thus skipping computationally demanding sequence alignments. For each protein sequence an MD5 hash digest is computed and looked up in a compact, embedded and read-only SQLite database. To reduce the risk of false identification due to hash collisions, amino acid sequence lengths of query and subject protein sequences are checked for equality. This combined procedure of identification via full-length protein sequence MD5 hash digests and related protein sequence length checks is subsequently referred to as alignment-free sequence identification (AFSI).

Remaining unidentified protein sequences are then searched against protein sequence clusters (PSCs), *i.e*. UniRef90 cluster representative sequences, using Diamond [29]. Alignment hits are filtered for mutual coverage and a sequence identity of at least 80% and 90%, respectively. Remaining alignment hits with sequence identities between 50% and 90% are then assigned to protein sequence cluster of clusters (PSCCs), *i.e*. UniRef50 clusters. Finally, preassigned annotations for identified and cluster-related protein sequences are looked up in the aforementioned SQLite database comprising gene symbols, protein products, dbxrefs to UniRef100, UniRef90 and UniRef50, RefSeq, clusters of orthologous (COG), enzyme commission (EC) categories and gene ontology (GO) terms [16,17,30–32].

To further improve the annotation of special interest genes, additional expert annotation tools are incorporated into the workflow allowing for fine grained annotation of closely related protein sequences that are indistinguishable by UniRef90 clusters alone. For instance, different alleles of antimicrobial resistance genes are annotated by AMRFinderPlus [33]. Furthermore, an integrated set of reference protein sequences with curated coverage and identity thresholds is used to refine annotations, thus allowing the standardized incorporation of external high-quality annotation resources, *e.g*. NCBI BlastRules and VFDB [16,34].

Finally, all gathered information is assessed to assign concluding annotations. CDS product names are amended and refined to follow protein nomenclature guidelines. CDSs without annotations are then (*i*) marked as hypothetical proteins; (*ii*) described by sequence-based characterizations, *i.e*. molecular weight and isoelectric point; (*iii*) screened for protein domains by HMMER using Pfam HMM profiles [27,35].

Results are provided in specification-compliant GFF3, EMBL and GenBank files. To foster streamlined submissions to member databases of the International Nucleotide Sequence Database Collaboration (INSDC), *e.g*. GenBank and ENA, further filtering and revision steps are implemented in an INSDC-compliance mode (`--compliant`) as some information-rich annotations are not fully-compatible with the strict validation rules of the INSDC. In addition, a compact human readable feature summary is presented in tabular file format. Genome and protein sequences are provided as Fasta files. Furthermore, sequences as well as characterizations and detected domains of hypothetical proteins are provided as Fasta and TSV files, respectively. To streamline automated downstream analysis and to encourage the incorporation of Bakta into larger analysis pipelines, all annotations and intermediate information are provided as detailed and standardized JSON files.

### Creation of a comprehensive taxon-independent database

Bakta takes advantage of a taxon-independent and comprehensive custom database integrating covariance models, HMMs, DNA and protein sequences. UPSs and protein cluster representative sequences of coding genes are enriched with pre-compiled information comprising gene symbols, protein products, EC numbers, COG and GO terms that are stored in a compact SQLite database. All database creation steps are automated as Bash and Python scripts and publicly available as part of the GitHub repository (https://github.com/oschwengers/bakta). The custom database is strictly versioned following a `<major>.<minor>` schema allowing for compatibility checks of the database schema as well as incremental minor updates.

For the annotation of non-coding features, bacterial covariance models for ribosomal RNA genes, ncRNA genes and ncRNA regulatory regions are downloaded and extracted from Rfam [21] via custom MySQL scripts and filtered by manually curated blocklists. DNA sequences of origins of replications and origins of transfer are downloaded and extracted from DoriC [24] and MOB-suite [36], respectively.

To annotate coding genes, binary MD5 hash digests of full-length amino acid sequences of all bacterial and phage related unique protein sequences from UniRef100 or UniParc [17], are computed and stored as UPSs along with protein sequence lengths and database identifiers, *i.e*. UniParc and UniRef100 to the SQLite database. In order to forestall potential hash collisions, the uniqueness of all computed MD5 hash digests is checked during this initial database creation step. Besides UniRef100 identifiers, gene symbols, protein products and related UniRef90 identifiers of taxonomically filtered UniRef100 records are stored as identical protein sequences (IPSs). Via this abstraction layer, UPSs of UniRef100 cluster members, *e.g*. unique sequence fragments, can be identified and linked to a common IPS. In addition, PSCs are created from UniRef90 records storing UniRef90 and related UniRef50 database identifiers, gene symbols and protein products. Likewise, PSCCs are created from UniRef50 records storing UniRef50 database identifiers and protein products. After this database initialization procedure, created UPSs, IPSs and PSCs records within the SQLite database are refined with annotations from external databases. Protein products and gene symbols are extracted from RefSeq non-redundant protein records [16], UniProt/SwissProt [17], AMRFinderPlus [33] or ISfinder [37]. Furthermore, annotations are enriched with additional information like EC numbers, COG functional categories and GO terms. All annotations are conducted and supersede each other according to the specificity of annotation sources. A more detailed description is provided in Supplemental Notes S1. Finally, PSCs which still remain unannotated, *i.e*. not being annotated with a protein product different from *hypothetical protein* or *uncharacterized protein*, are subsequently scanned against Pfam protein family HMMs [35] and annotated upon sufficient hits accordingly. The import of UPS, IPS, PSC and PSCC records and all conducted annotations are logged for the sake of transparency, enabling potential later inspections. This log file is hosted at Zenodo along with the database itself. To reduce the total size of the SQLite database, prefixes are removed from all internal and external database identifiers. This procedure is reversed at runtime to reproduce original database identifiers. Finally, the SQLite database is defragmented and reduced in size by the SQLite `VACUUM` pragma.

For the integration of high-quality annotation sources from external databases that are available at runtime, a general protein sequence-based expert annotation system is compiled. Therefore, protein sequences, gene symbols, protein products, query and subject coverage thresholds, sequence identity thresholds and priority ranks are stored for protein sequences from VFDB [34] and NCBI BlastRules [16]. More information is provided in Supplemental Notes S2.

The deeper analysis of hypothetical proteins is a distinct task in Bakta’s annotation workflow. Therefore, Pfam [35] HMMs of types different from *family* are downloaded and included in the database for the detection of conserved sequence domains within these proteins of unknown functions at runtime.

## Results

### Comparison of annotated features

To illustrate and compare all aspects of Bakta’s functionality we evaluated its performance and benchmarked it against other software tools. For these comparisons, we focused on state-of-the-art command line annotation command line software tools providing likewise short wall clock runtimes and low resource consumptions, *i.e*. Prokka and DFAST. For the sake of comparability, we chose the genome of *Escherichia coli* O26: H11 str. 11368 (GCF_000091005.1) that was also used by the authors of DFAST and annotated this genome with Prokka 1.14.6 [6], DFAST 1.2.11 [7] and Bakta 1.1. To additionally provide a preliminary comparison with annotation tools implementing a more elaborated but also more computationally demanding workflow, we complemented this set with the latest RefSeq annotation annotated with PGAP 5.2 [16]. A detailed comparison comprising distinct numbers of predicted, identified, functionally annotated and database cross-referenced CDS as well as numbers of predicted and annotated further feature types is summarized in Table 1.

**Table 1.** Comparison of annotation results of *Escherichia coli* O26: H11 str. 11368

| Feature types | PGAP | Prokka | DFAST | Bakta |
| --- | --- | --- | --- | --- |
| Total CDS | 5,794 | 5,754 | 5,740 | 5,841 |
| Identified proteins | 5,550 | — | — | **5,738** |
| With COG | — | 113 | **3,952** | 3,277 |
| With EC | 1,518 | 1,042 | 1,217 | **1,562** |
| With GO | — | — | — | **3,474** |
| Unknown function <sup>a</sup> | 423 | 1,808 | 1,358 | **225** |
| sORF <sup>b</sup> | 44 | — | — | **82** |
| Total tRNA | 102 | 106 | 106 | 106 |
| tRNA | 101 | 105 | 105 | 105 |
| tmRNA | 1 | 1 | 1 | 1 |
| Pseudo | — | — | — | 3 |
| rRNA | 22 | 22 | 22 | 22 |
| Total ncRNA | 17 | **295** | — | 289 |
| Genes | 10 | — | — | **223** |
| Regulatory regions | 6 | — | — | **66** |
| Miscellaneous |  |  |  |  |
| CRISPR array | 2 | 2 | 2 | 2 |
| Origins of replication | — | — | — | **4** |
| Computational resources |  |  |  |  |
| Wall clock runtime [mm:ss] <sup>c</sup> | — | 4:13 | **3:48** | 7:09 |
| RAM [GB] | — | **1.2** | 1.8 | 4.4 |

| DB size [GB] | — | **0.6** | 3.3 | 53 |
| --- | --- | --- | --- | --- |

Note: Numbers represent annotated features. Prokka, DFAST and Bakta were executed with default parameters providing all relevant information, *e.g.* genus and species, assembly status and sequence topology, a detailed list of command lines is provided in Supplemental Notes S3; annotations from PGAP were downloaded in GenBank file format from RefSeq.
<sup>a</sup> Protein sequences of unknown function; product denoted as *hypothetical protein*, *putative protein*, *uncharacterized protein* or *conserved predicted protein*.
<sup>b</sup> sORFs shorter than 29 amino acids
<sup>c</sup> Best out of three wall clock runtimes executed with 8 threads.

First, we compared the regional prediction of various features including coding, non-coding and further genomic features. Regarding tRNAs, tmRNAs, rRNAs and CRISPR arrays all tools predicted equal or comparable numbers of features. Prokka annotated the highest total number of ncRNAs whereas only PGAP and Bakta were able to distinguish between ncRNA genes and ncRNA regulatory regions. Taking this into account, Bakta predicted the highest number of ncRNA genes (n=223) and regulatory regions (n=66). Moreover, Bakta was the only tool predicting origins of replication (n=4). Regarding CDSs, Bakta (n=5,841) and PGAP (n=5,794) predicted more genes than Prokka (n=5,754) and DFAST (n=5,740) which we attribute largely to the detection of small proteins by Bakta (n=82) and PGAP (n=44) that are not predicted *de novo* by Prodigal [15] and MetaGeneAnnotator [14] used by Prokka and DFAST, respectively.

Second, we compared the identification and functional annotation of predicted and detected CDSs. In contrast to Prokka and DFAST, Bakta (n=5,738) and PGAP (n=5,550) were able to precisely identify publicly-known protein sequences and to assign stable database identifiers referring to RefSeq [4] and UniRef100 [17]. In terms of functional CDSs annotation, Bakta and PGAP provided more functional descriptions resulting in notably fewer CDSs annotated as *hypothetical protein*. To further assess the annotation performance of CDSs, we also looked for the assignment of functional ontologies, *i.e*. COG, EC numbers and GO terms, as these are valuable resources for downstream analysis. Here, DFAST assigned the highest number of COG identifiers (n=3,952), closely followed by Bakta (n=3,277). Bakta (n=1,562) assigned the most EC numbers, followed by PGAP (n=1,518), DFAST (n=1,217) and Prokka (n=1,042). In this benchmark, Bakta was the only tool that assigned GO terms (n=3,474).

### Runtime performance and resource consumption

Resource consumption and runtime characteristics of software tools are important factors for the annotation of bacterial genomes conducted on either local consumer hardware providing only limited computing resources or larger server machines within high performance or cloud computing infrastructures providing scalable computing resources. To compare aforementioned tools and to provide guidance on resource consumptions for different scenarios, we compared wall clock runtimes, memory consumptions and storage requirements. Therefore, we executed and monitored all tools three consecutive times on a server machine with 4 Intel Xeon E5-4627 CPUs and 40 cores in total. However, to provide exemplary runtimes that are also achievable on consumer hardware, we restricted available CPU resources to 8 threads. Results for the best out of three executions are provided in Table 1. Wall clock runtimes of Prokka (4:13 m:ss) and DFAST (3:48 m:ss) were considerably shorter than those of Bakta (7:09 m:ss). Likewise, Prokka (1.2 GB) and DFAST (1.8 GB) required less memory than Bakta (4.4 GB). Also, database sizes of Prokka (0.6 GB) and DFAST (3.3 GB) are considerably smaller than that of Bakta (53 GB). However, at the time of writing, Bakta’s underlying database v3.0 comprises more than 90.5 million UniRef90 reference sequences and hash digests of more than 214.8 million unique protein sequences from UniParc and UniRef100. Hence, the numbers of protein sequences contained in the databases of Prokka (n=32,148) and DFAST (n= 405,076) are exceeded by several orders of magnitude. Taking this into account, the performance of the Bakta pipeline can be seen as a huge relative speedup, as it offers a big increase in depth of analysis compared to Prokka and DFAST solely at the cost of a very moderate increase in wall clock runtimes. This acceleration was achieved via the AFSI approach that drastically reduced the number of required CDS alignments to 110 in this benchmark. Wall clock runtimes required to conduct homology searches for these remaining protein sequences are further reduced by using Diamond [29] using its new fast mode. Hence, even though Bakta provides a much larger and more comprehensive annotation database, it is able to annotate bacterial genomes within wall clock runtimes roughly comparable to Prokka and DFAST even on standard consumer hardware.

To assess both the vertical scalability of each tool and the effects of AFSIs on overall runtime performances, we conducted a second benchmark measuring wall clock runtimes using varying numbers of CPU cores. Therefore, we created a Bakta version with deactivated AFSI logic which is subsequently referred to as Bakta w/o AFSI. In this experiment, DFAST consistently provided the shortest runtimes within each bin of available CPU cores followed by Prokka and Bakta (Fig. 2) in line with wall clock runtimes of the first benchmark. Each tool exhibited a solid scalability between 1 and 16 CPU cores. The addition of further CPU cores contributed only neglectable runtime reductions. Furthermore, Bakta consistently showed considerably reduced wall clock runtimes compared to Bakta w/o AFSI for all measured numbers of CPU cores demonstrating the acceleration benefits of AFSIs. However, it must be noted that the annotated *E. coli* genome is part of RefSeq and thus Bakta’s database comprises a large proportion of these protein sequences. To assess potential AFSI accelerations for species that are not contained in the public databases, we repeated this experiment with a hitherto unknown genome of a recently described new *Pseudocitrobacter* species [38]. Therefore, raw sequencing reads were downloaded from ENA (ERR3255970), quality filtered with fastp (0.20.1) [39] and assembled with Unicycler (0.4.8) [40]. In line with our expectations that the power of AFSI runtime accelerations depends on the number of identifiable protein sequences which in turn roughly correlates with the taxonomical proximity of the species at hand with those comprised by the public databases, AFSIs showed only moderate runtime advantages in this experiment (Supplemental Fig. S1).

**Figure 2:**
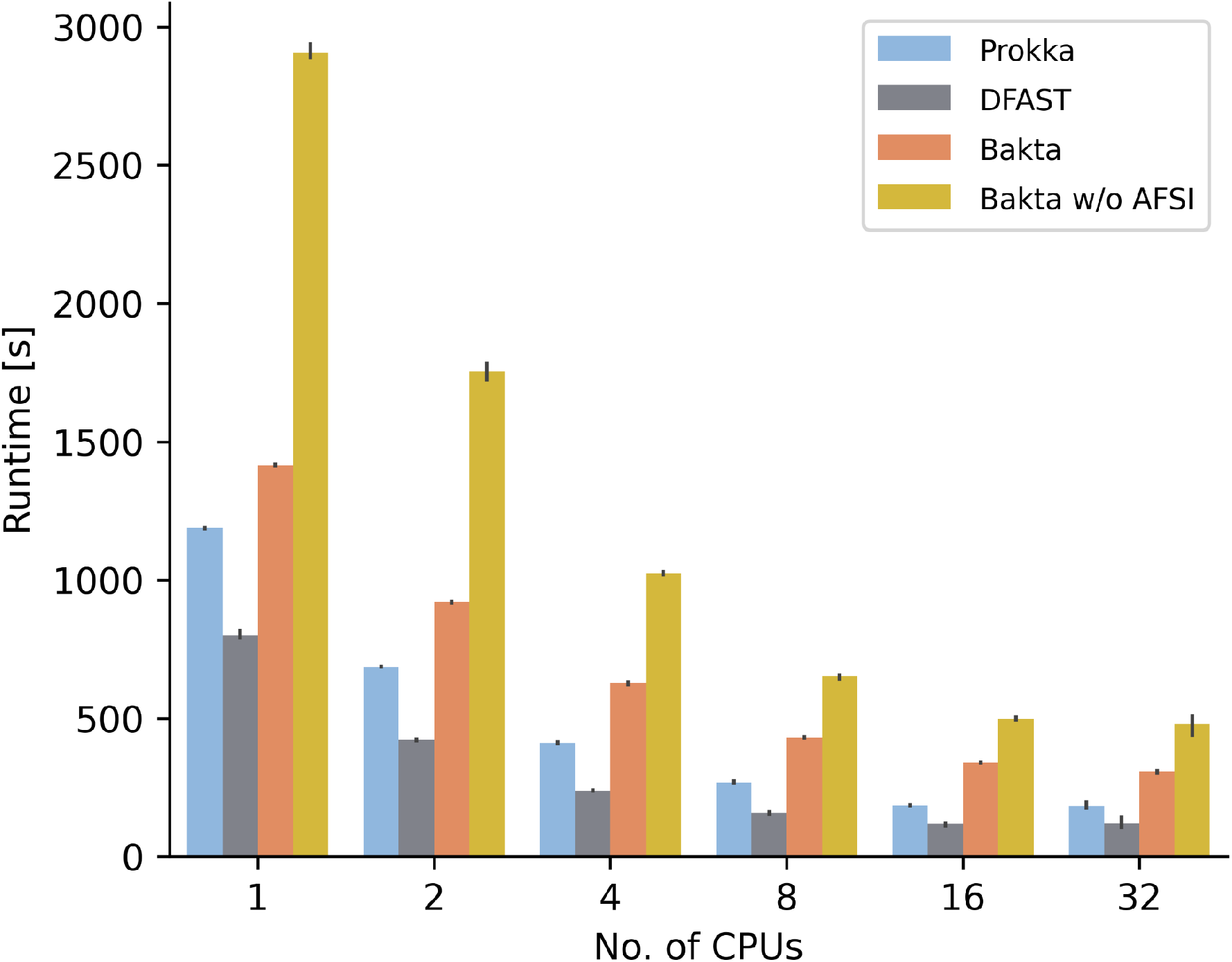
Comparison of wall clock runtimes. Runtimes of Prokka, DFAST, Bakta and Bakta w/o AFSI annotating *Escherichia coli* O26: H11 str. 11368 were measured three consecutive times using varying numbers of CPUs on a server machine with 4 Intel Xeon E5-4627 CPUs and 40 cores in total.

### Functional annotation performance benchmark

We envision Bakta as a suitable alternative to existing command line annotation software tools, *e.g*. Prokka and DFAST. Furthermore, we see great potential for integration into larger high-throughput analysis pipelines, *e.g*. Tormes [8], ASA^3^P [9], Bactopia [10] and Nullarbor [11], enabling taxonomically untargeted workflows. Hence, we compared the functional annotation performance of Bakta against aforementioned tools over a broad taxonomic range of species. Therefore, we counted numbers of predicted CDSs and those annotated as *hypothetical protein* in total and genome-wise manner. Moreover, we counted the numbers of identified protein sequences and detected small proteins by Bakta.

In a first experiment we annotated 35 taxonomically diverse bacterial genomes from RefSeq [4]. This benchmark dataset comprises many bacterial pathogens, *e.g*. ESKAPE species, as well as commensal and environmental species. RefSeq assembly accessions and detailed benchmark results for all genomes are available in Supplementary Table S1. In addition, Bakta result files for each test genome are publicly hosted at Zenodo (DOI: <u>10.5281/zenodo.5253552</u>) to serve as annotation examples. In this benchmark, DFAST predicted the fewest CDSs (n=127,053) followed by Prokka (n=130,360) and Bakta (n=130,683). As both Prokka and Bakta internally use Prodigal for the *de novo* prediction of CDSs, the difference of 323 predicted CDSs is mainly due to the detection of 235 small proteins by Bakta as well as to differences in the internal feature overlap filters of both tools. Regarding the functional annotation, Bakta achieved a total ratio of CDSs annotated as *hypothetical protein* as low as 10.6% (n=13,902) outperforming DFAST (n=40,128) and Prokka (n=53,656) which achieved total ratios of 31.6% and 41.2%, respectively (Fig. 3). Within the set of benchmarked tools, only Bakta was able to identify publicly known unique protein sequences. 94.2% (n=123,105) of all predicted CDSs (n=130,683) were precisely identified via AFSI. The genome-wise minimum and maximum ratios of identified protein sequences reached 77.2% and 99.9%, respectively. These results show that a large proportion of CDSs can be identified via AFSIs over a broad and diverse taxonomic range of genomes thus facilitating the assignment of public identifiers. It goes without saying that these identified CDSs also comprised proteins of unknown functions, *i.e*. annotated as *hypothetical protein*. However, in these particular cases, assigned public identifiers are of even higher value as they support further investigations taking into account additional information from external databases.

**Figure 3:**
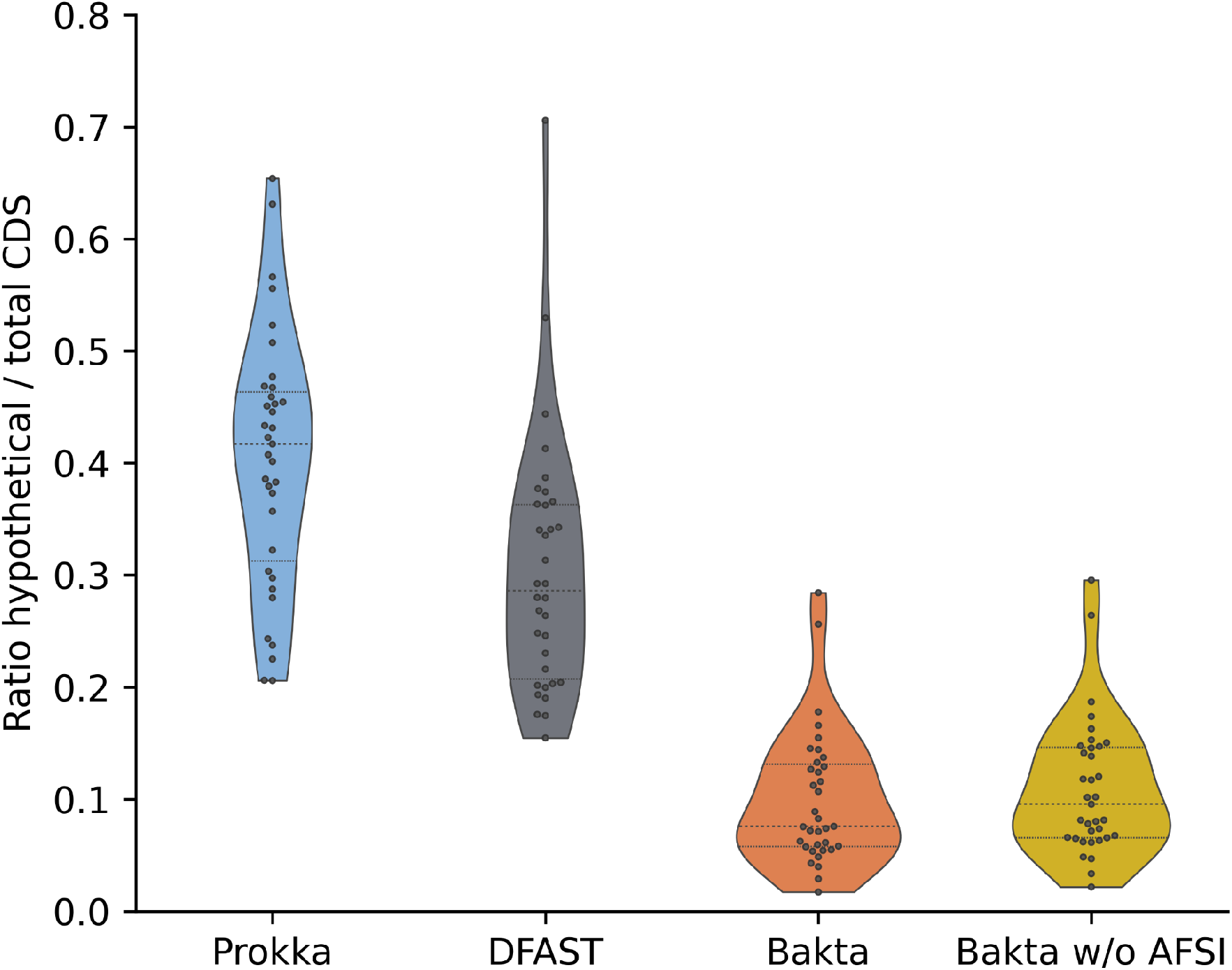
Proportion of protein sequences annotated as *hypothetical protein*. Distributions of genome-wise ratios of numbers of total CDS and those annotated as *hypothetical protein* are shown for 35 selected RefSeq genomes comprising species of high medical and biotechnological relevance.

One limitation of this set of RefSeq benchmark genomes is the fact that all of these genomes are contained within RefSeq and thus are also contained in Bakta’s custom database. Therefore, it is not surprising that most of these protein sequences could be identified and functionally annotated via AFSI. However, in order to show that wall clock runtime acceleration in conjunction with the assignment of database identifiers constitute the main advantage of the AFSI approach rather than the improvement of annotation qualities, we benchmarked the functional annotation performance of Bakta w/o AFSI. The internal workflow of this version defaults to mere homology searches, *i.e*. Diamond sequence alignments against PSC sequences, without conducting any AFSIs at all. In this benchmark, Bakta w/o AFSI achieved a total ratio of CDSs annotated as *hypothetical protein* of 11.5% (n=15,066) resulting in a difference between Bakta with and without AFSI as low as 0.9% (n=1,164). Hence, we conclude that AFSIs make only small contributions to the functional annotation of protein sequences. However, it provides huge potential to avoid computationally expensive sequence alignments and profile searches besides the precise identification of publicly known protein sequences.

To address the discussed limitations of the RefSeq benchmark dataset we ran a second experiment to assess the functional annotation performance on a large set of genomes that are not covered by those public databases that are used within the database build procedure. Therefore, we screened the GenBank database for genomes meeting the following criteria: (*i*) they have a strain designation to exclude metagenome-derived genomes; (*ii*) they have explicitly been excluded from RefSeq due to an undefined genus; (*iii*) they do not miss certain features, *e.g*. tRNA and rRNA. The resulting 362 genomes (Supplementary Table S2) were annotated with Prokka, DFAST and Bakta without providing any taxonomic information. In this benchmark, DFAST predicted the fewest CDSs (n=1,113,906) followed by Bakta (n=1,127,661) and Prokka (n=1,128,187). On average, Bakta achieved a total ratio of CDSs annotated as *hypothetical protein* as low as 25.4% (n=286,406) outperforming DFAST (n=457,245) and Prokka (n=548,167) which achieved total ratios of 41.1% and 48.6%, respectively. Figure 4 shows the distributions of genome-wise hypothetical protein proportions. Even though none of these genomes are contained in RefSeq, Bakta identified 26.6% (n=299,410) of all CDSs via AFSI. However, this time genome-wise minimum and maximum ratios of identified protein sequences ranged between 0% and 99.9%, respectively with a median of 10.4%.

**Figure 4:**
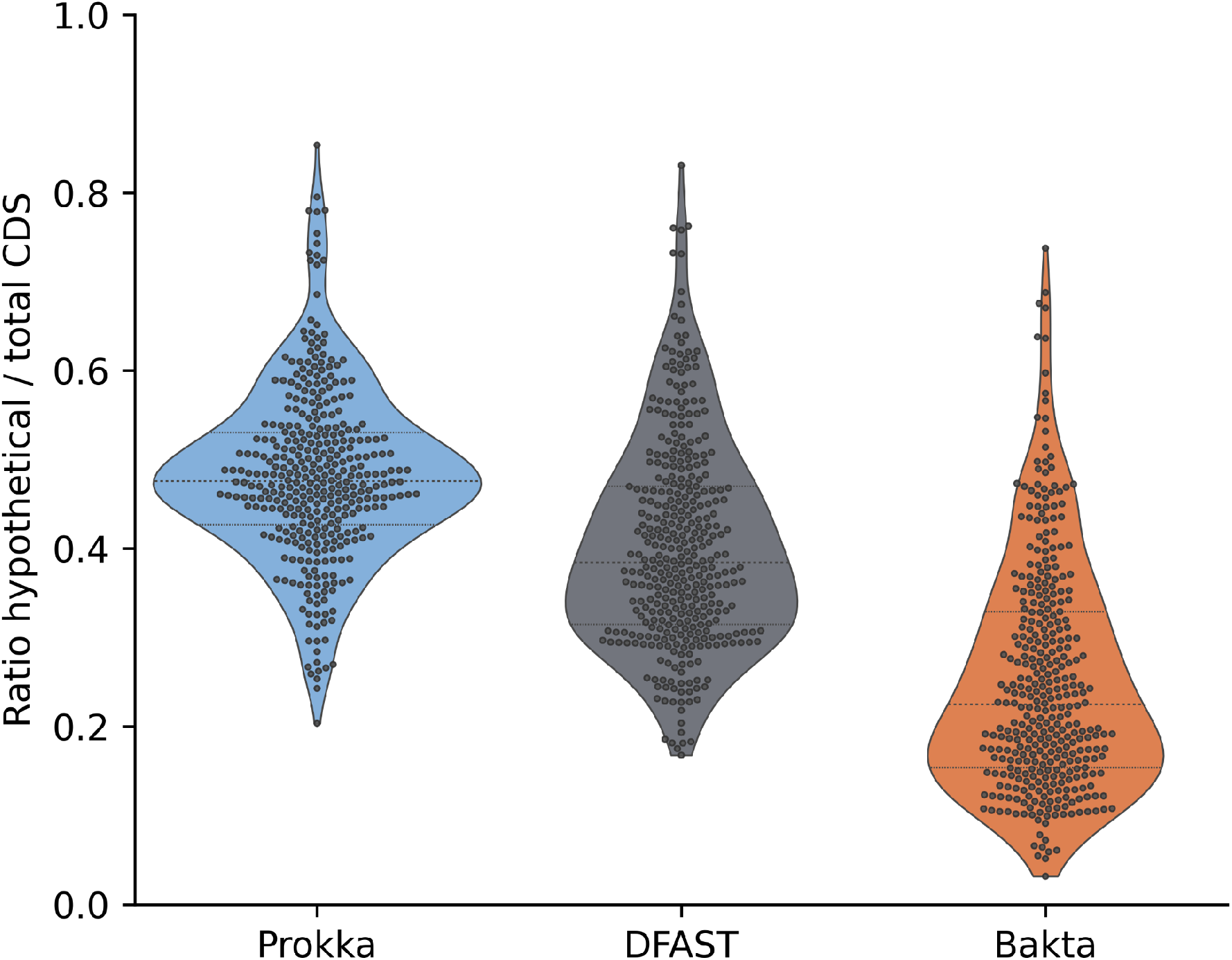
Proportion of protein sequences annotated as *hypothetical protein*. Distributions of genome-wise ratios of numbers of hypothetical proteins and total CDS are shown for 362 GenBank genomes comprising species of undefined genera.

As small proteins are known to play important roles in many processes, *e.g*. regulation [41], virulence [42,43] and sporulation [44], we investigated the functional descriptions of all detected small proteins from the RefSeq benchmark experiment in order to assess the relevance and impact of their annotation. Table 2 summarises the numbers of detected small proteins aggregated by key words contained in the proteins’ product descriptions. These results indicate that the small proteins detected by Bakta in this benchmark are involved in a broad range of important processes of high relevance to pathogenicity as well as more general cellular house-keeping processes.

**Table 2.** Functional categories of small proteins detected by Bakta.

| Function <sup>a</sup> | Number of detected small proteins |
| --- | --- |
| attenuator and leader peptides | 53 |
| membrane | 10 |
| phage | 8 |
| regulation | 7 |
| phenol-soluble modulin | 7 |
| toxin-antitoxin systems | 6 |
| toxins | 5 |
| sporulation | 5 |

<sup>a</sup> Extracted from annotated product descriptions.

### Annotation of metagenome-assembled genomes

Since 2004, advances in the sequencing of entire microbial communities comprising uncultivated organisms combined with new bioinformatics methodologies [45] revealed hitherto unknown taxa and led to a burst of new bacterial genomes [46–49]. However, as large proportions of the proteins encoded by these genomes are of unknown functions, the automated annotation of these genomes remains challenging. To address these issues, we complemented the annotation workflow of Bakta with a fallback stage to further expand the recognizable sequence space. Protein sequences that cannot be identified neither by IPSs nor PSCs are annotated by PSCCs, *i.e*. UniRef50 clusters. To assess the annotation performance of Bakta and to compare it against Prokka and DFAST, we compiled a benchmark set of high-quality MAGs. Therefore, we screened 7,903 published MAGs [46] that have been assembled from more than 1,500 public metagenomes meeting the following criteria: (*i*) a CheckM [50] complete score larger than or equal to 95.0; (*ii*) a CheckM contamination score smaller than or equal to 1.0; (*iii*) a taxonomical assignment within the bacterial GTDB lineage. Using this benchmark dataset comprising 198 MAGs (Supplementary Table S3) covering a diverse taxonomic range (Supplemental Fig. S2), Bakta achieved on average a total ratio of CDSs annotated as *hypothetical protein* as low as 24.2% (n=138,282) outperforming DFAST (n=232,516) and Prokka (n=279,352) which achieved total ratios of 41.3% and 49.0%, respectively. Figure 5 shows the distribution of genome-wise *hypothetical protein* ratios. For 46.5% (n=92) of all MAGs Baka achieved the lowest genome-wise *hypothetical protein* ratio. Interestingly, even in this metagenomic setup, Bakta was able to precisely identify 38.6% (n=220,753) of all predicted CDSs (n=572,213) via AFSI.

**Figure 5:**
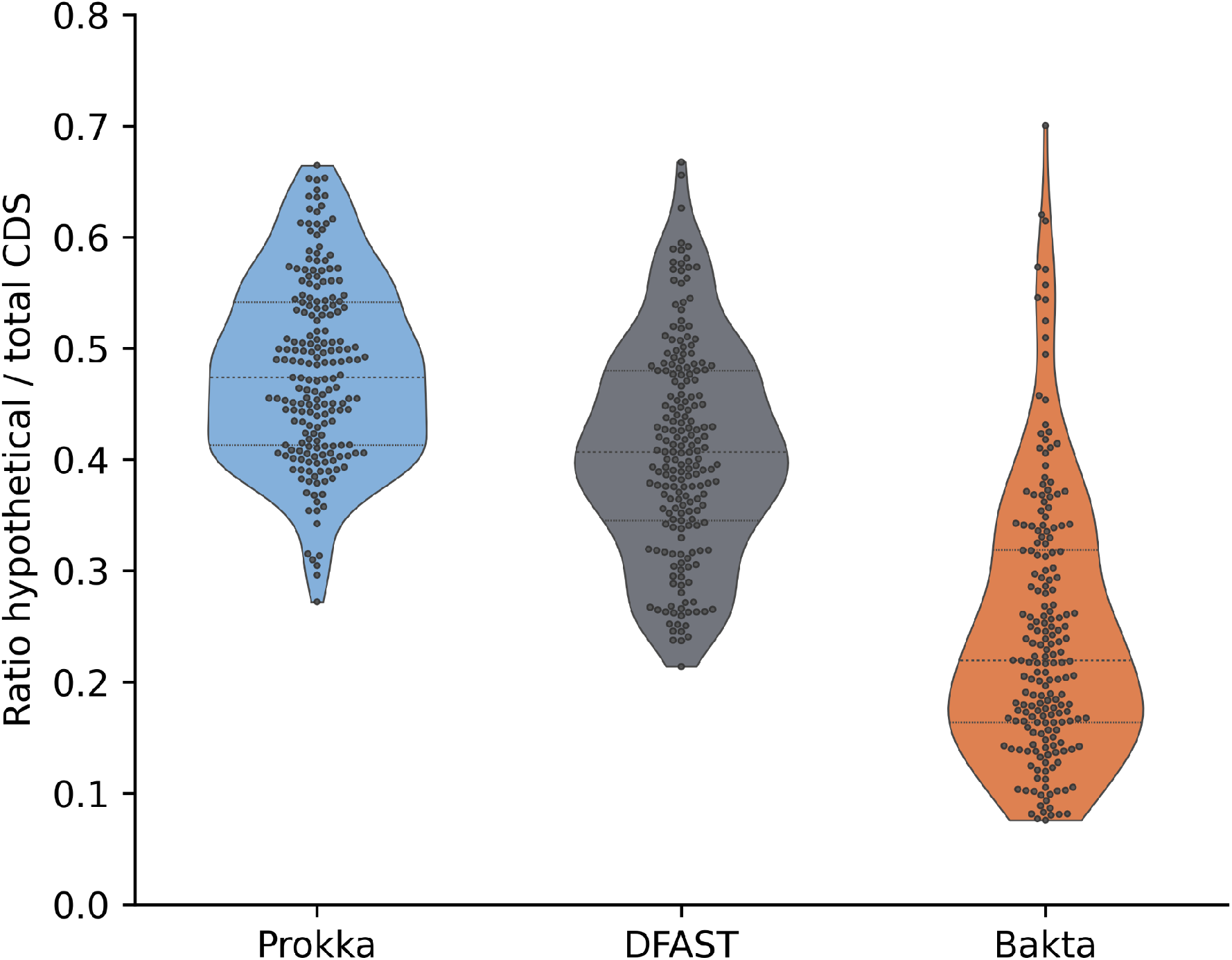
Proportion of protein sequences annotated as *hypothetical protein*. Distributions of genome-wise ratios of numbers of total CDS and those annotated as *hypothetical protein* are shown for 198 bacterial high-quality metagenome-assembled genomes screened by genome completeness and contaminations.

### INSDC-compliant annotation results

The INSDC is a long-standing initiative synchronizing the major public DNA sequence databases DDBJ, ENA and GenBank. The submission of annotated genomes to these databases is a prerequisite for the publication of genomic data in most scientific journals. Hence, the INSDC-compliant export of annotation results in INSDC flat file formats is a crucial task of contemporary genome annotation pipelines. To streamline this process and to provide information-rich annotations compatible with the strict validation rules of the INSDC, we implemented a compliance mode that can be used via the `--compliant` option. To assure INSDC compliance of Bakta’s output files, we validated the EMBL flat files created for the 35 genomes of our benchmarking dataset (Supplementary Table S1) using the Webin-CLI submission tool version (4.0.0) provided by the ENA [51]. All tested files were successfully validated without errors or warnings. In addition, annotated genomes can be submitted to GenBank via NCBI’s `table2asn_GFF` tool using Bakta’s GFF3 and Fasta files.

### Convenient and scalable web-based annotations

Command line software tools are essential for the timely analysis of large bacterial cohorts. They facilitate rapid and scalable annotations conducted either on local hardware or within cloud computing infrastructures, respectively. Nevertheless, graphical user interfaces are sometimes favored due to supplemental features, as for instance the interactive visualization of results. To additionally address these demands and to ease the access to the software for users without sufficient command line experience, we developed an accompanying and convenient web application driven by a scalable cloud-based backend (Supplementary Notes S4) available at https://bakta.computational.bio.

This web application provides an interactive GUI wizard that supports the user in the upload of input data, the specification of related metadata as well as the configuration and submission of annotation jobs (Fig. 6). For instance, it automatically parses the uploaded genome in Fasta file format [52] and provides a replicon table widget that aids the user with the provision of precise metadata for each replicon sequence within the genome. Furthermore, the configuration of annotation parameters is supported via a taxon autocompletion mechanism for genus and species information that takes advantage of the ENA Taxonomy REST API [51]. Finally, annotation results are provided in various manners. Firstly, a set of aggregated feature counts provides a broad picture of the genome. Secondly, a searchable data table contains detailed information on each predicted feature providing a fulltext search and filter capabilities. To allow deeper investigations of certain genes taking into account additional external information, listed features are linked to related public database records via assigned dbxrefs. Last but not least, an interactive visualization of the annotated genome is provided via an igv.js [53] based genome browser. CDSs features are colored according to COG functional categories.

**Figure 6:**
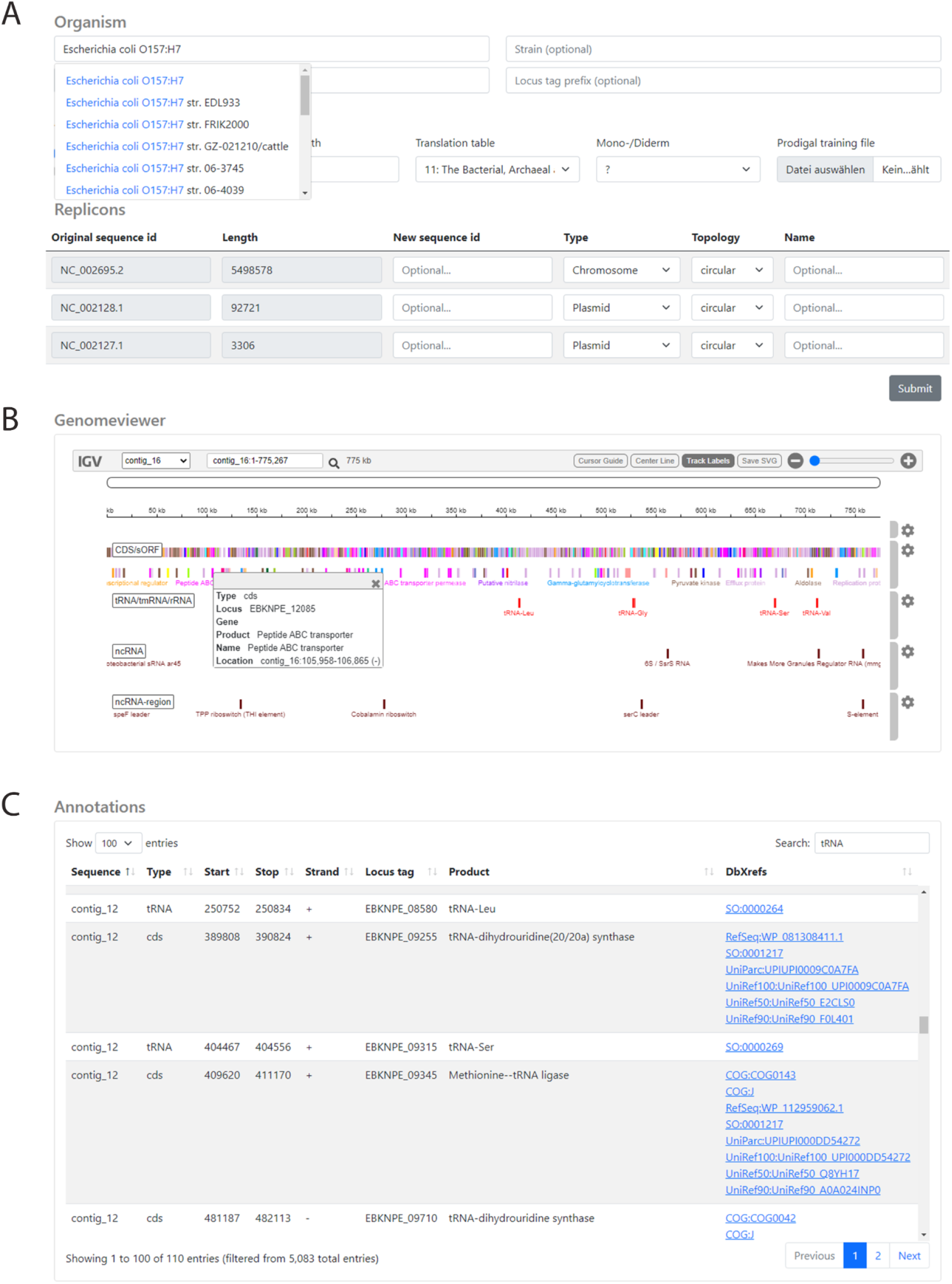
GUI screenshots of the Bakta web version. (A) Submission page with metadata input fields providing taxon autocompletion support for genus and species (top) and replicon table editor (bottom). (B) An igv.js-based genome browser visualizing annotated features. CDS-features are colored according to the annotated COG functional category. (C) Interactive annotation table providing search and filter features. Annotated dbxrefs are linked to target databases.

We would like to emphasize that this web application can also be used to visualize offline annotation results conducted by using the command line version. Therefore, the web application provides an offline viewer accepting Bakta’s JSON result files which are parsed and visualized locally within the browser without sending any data to the server.

## Discussion

The progress of DNA sequencing technologies in recent years has led to tremendously increasing numbers of bacterial genome sequences. In turn, the implied huge increase in computational workloads has driven the development of rapid and lightweight command line annotation pipelines as suitable offline alternatives to established online annotation services. These tools achieve very short wall clock runtimes and support additional user-provided annotation sources. However, this is achieved at the cost of smaller database sizes and results in less standardized annotation workflows. To address these issues, we developed Bakta, a new command line annotation software tool aiming at a well-balanced tradeoff between runtime performance and comprehensive annotations. This new software tool is implemented in Python 3 and can be installed on any UNIX system via Conda, Docker and Singularity. It scales to multiple cores and allows the annotation of a typical genome within approximately 10 minutes.

In contrast to existing light-weight annotation software tools, Bakta also detects and annotates sORFs of small proteins. Two decades ago, the existence of many of these small proteins was experimentally verified expanding the prokaryotic genomic repertoire. Existing lightweight command line annotation tools fail to detect these small proteins through using contemporary *de novo* gene prediction tools [14,15] alone. To the best of our knowledge, Bakta is currently the only lightweight annotation software tool that is able to detect and annotate these small proteins. However, it must be stated that Bakta is not able to predict these small protein coding genes *de novo* either. Instead, it identifies known sORF protein sequences via AFSI and additionally conducts very strict homology searches to find and annotate these sequences. Thus, Bakta helps to shed light on these otherwise genomic blind spots. This approach however has an obvious drawback as it is not able to predict hitherto unknown sORFs. Hence, the integration of dedicated sORF prediction tools [54,55] into this workflow might help to improve on this issue.

Existing lightweight annotation software tools accelerate the execution of their workflow by using hierarchical or taxonomically targeted annotation databases. In contrast, Bakta provides a single taxonomically untargeted database. By doing so, it facilitates the integration into larger high-throughput analysis pipelines that might be executed in a taxon-independent manner. Also, it allows the annotation of rare bacterial species for which no or only few high-quality reference genomes exist. It goes without saying that larger and more comprehensive databases have negative effects on overall wall clock runtimes. To mitigate this effect and nevertheless keep runtimes as low as possible, Bakta follows a different approach: the reduction of required sequence alignments. Therefore, we introduce AFSI as a new approach to this issue. To the best of our knowledge, this has not been used before in the context of protein sequence identifications and genome annotation. We demonstrated that this approach is capable of identifying large proportions of coding genes on large sets of taxonomically diverse genomes. Hence, numerous computationally expensive homology searches can be avoided and thus the overall annotation process is massively accelerated. Interestingly, we could demonstrate that AFSI also performs well on MAGs of potentially unknown species. The precise identification of protein sequences via AFSI has various advantages besides mere wall clock runtime reductions. It furthermore provides a valuable tool for the tracing of certain genes, *e.g*. antimicrobial resistance genes within populations or during outbreaks. Furthermore, it facilitates streamlined comparative analysis and compliance with FAIR [56] data principles by cross linking genome features to public database records, *e.g*. RefSeq [16] and UniRef [17]. Often, these databases are in turn linked to other databases that additionally contribute to a more comprehensive and sophisticated picture of these genomic sequences. Especially for protein sequences of unknown functions, *i.e*. proteins annotated as *hypothetical protein*, the interconnection of database records provides a helpful tool for further investigations.

An important aspect that must not be overlooked are potential hash collisions which might lead to false identifications and hence wrong annotations. In its current version 1.1 Bakta uses the MD5 hash algorithm due to its fast computation and short hash sum length. So far, no hash collisions could be detected during the database creation procedures incorporating more than 214.8 million distinct protein sequences. Of course, this cannot be assured for future database releases comprising an expanded protein sequence space and it might become necessary to switch to other hashing algorithms, *e.g*. SHA256. This might additionally increase future database sizes. To accelerate the lookup of large numbers of these hash digests, they are stored in a compact binary format within an SQLite database. It should be noted that this might have severe negative effects on the overall runtime performance if the custom database is stored on network attached storage volumes - a common situation on high-performance compute clusters and cloud computing infrastructures. For these setups, we highly recommend using a local copy of the database.

The precise annotation of CDSs conducted by Bakta is based on alignment-free detections of IPSs complemented by alignment-based homology searches for PSC homologues. However,depending on taxonomic distributions and evolutionary selection pressures, sequence conservation of protein family members may vary significantly. Hence, the AFSI of certain protein sequences belonging to more heterogeneous protein families might not always be possible. Likewise, appropriately precise annotations of CDSs belonging to closely related but nevertheless distinct protein families might not be achievable via PSCs. To facilitate more precise annotations of these CDSs, Bakta complements its annotation workflow by taking advantage of so-called expert annotation systems. At the time of writing two expert annotation systems are implemented: one to specifically target antimicrobial resistance genes and a general protein sequence-based system integrating multiple external high-quality annotation sources. The expansion of these expert systems are subject for further improvements.

The recent progress in metagenomics nowadays allows the sequencing of entire microbial communities and to reconstruct MAGs *in silico* thus providing access to hitherto unknown genomes of unculturable organisms. The annotation of these genomes is key to many downstream analyses, such as metabolic pathway predictions. However, the annotation of these genomes via reference genomes or taxonomically targeted databases becomes difficult or even impossible for rare or unknown species that are covered poorly or not at all by public databases. To improve the annotation of these genomes we implemented an additional annotation step. We demonstrated that Bakta is able to annotate large proportions of many MAGs’ protein sequences and outperforms other annotation software tools.

In conclusion, we have developed the new command line software tool Bakta, and we demonstrated that it improves on existing rapid annotation tools for bacterial genomes in various ways: (*i*) Bakta outperforms existing tools in terms of functional annotation of CDSs over a broad taxonomic range of both known and unknown species; (*ii*) Bakta is able to detect and annotate small proteins which are not predicted by contemporary *de novo* gene prediction tools, as for instance Prodigal [15] and MetaGeneAnnotator [14]; (*iii*) Bakta precisely identifies publicly known protein sequences and assigns stable database identifiers from RefSeq [16] and UniProt [17]; (*iv*) Bakta’s functional annotation workflow is accelerated by a new AFSI approach; (*v*) Bakta takes advantage of sequence metadata to improve the structural prediction of CDSs; (*vi*) Bakta provides equivalent or more comprehensive annotations of CDSs with functional categories, *i.e*. COG, EC numbers and GO terms. Therefore, we consider Bakta as a useful and valuable novel tool for the comprehensive and timely annotation of bacterial genomes, even on standard consumer hardware. In addition, we have developed a user-friendly web version providing interactive visualizations taking advantage of a highly-scalable cloud based backend.

## Acknowledgements

We gratefully thank Karina Brinkrolf for comments and valuable feedback and Anna Rehm and Julian Hahnfeld for code contributions.

