## Supplementals for "Bakta: Rapid & standardized annotation of bacterial genomes via alignment-free sequence identification"

Affiliation:

Keywords:

GitHub: https://github.com/oschwengers/bakta

Zenodo: DOI 10.5281/zenodo.4247252

Web: https://bakta.computational.bio

### Supplemental Figures

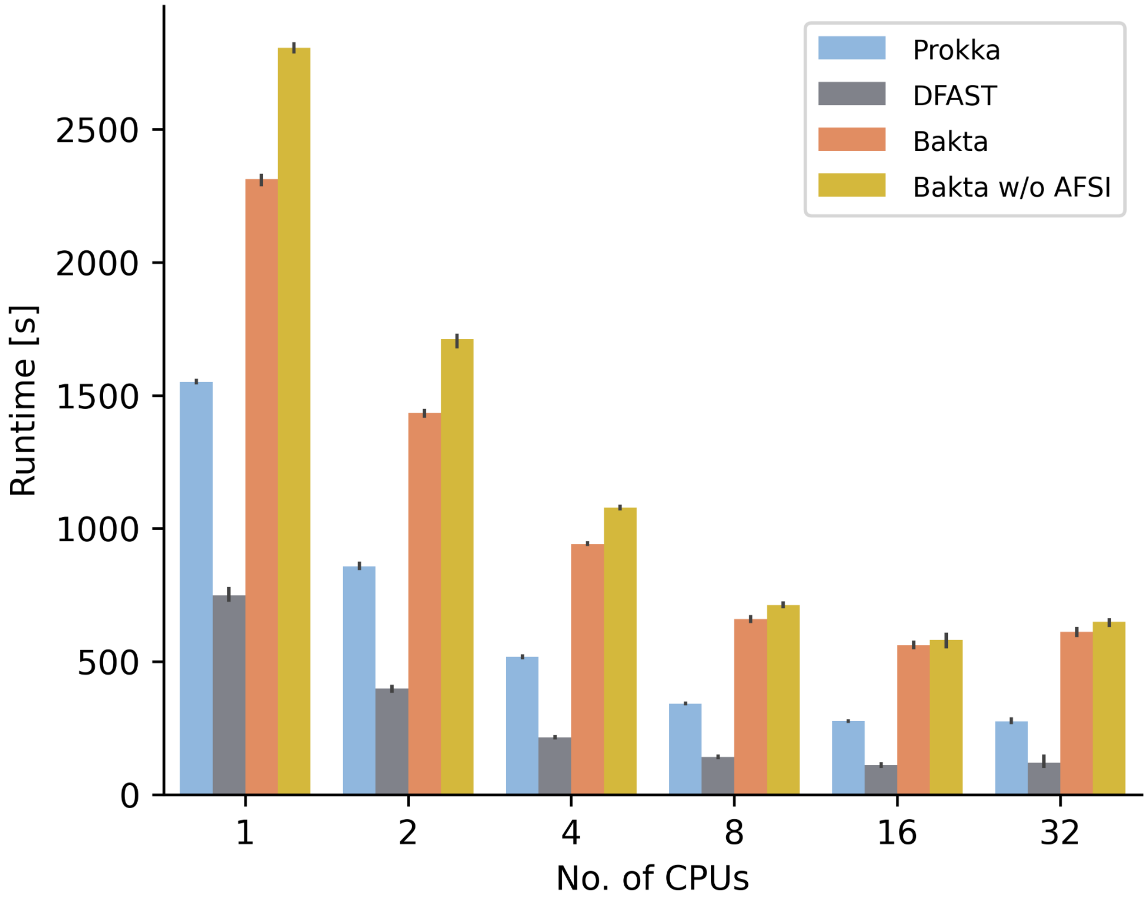


**Supplemental Figure S1.** Comparison of wall clock runtimes. Runtimes of Prokka, DFAST, Bakta and Bakta w/o AFSI annotating a *Pseudocitrobacter* genome were measured three consecutive times using varying numbers of CPUs on a server machine with 4 Intel Xeon E5-4627 CPUs and 40 cores in total.

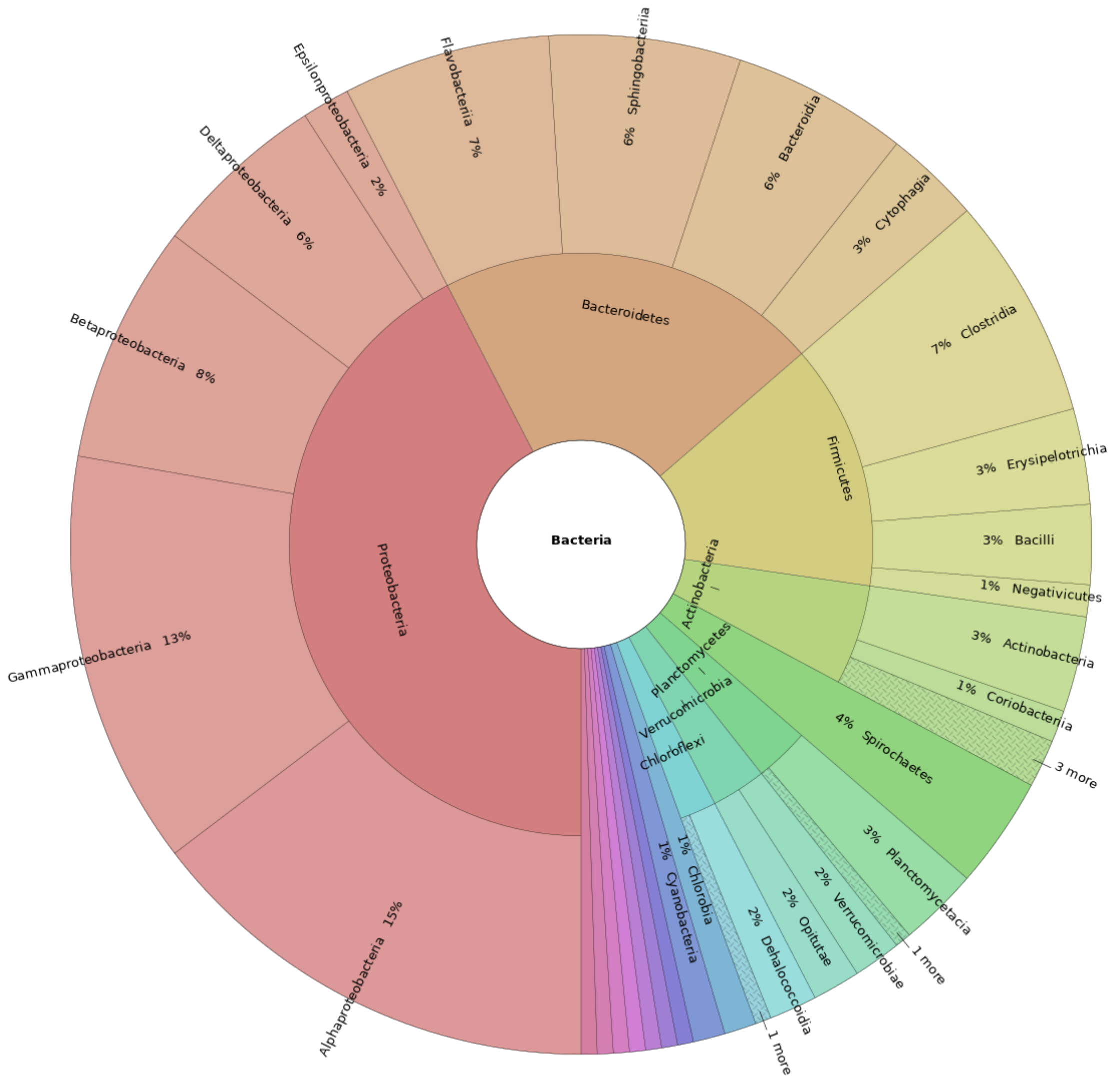


**Supplemental Figure S2.** Taxonomic distribution of metagenomic-assembled benchmark genomes. 198 bacterial MAGs were selected from 7,903 published MAGs [1] by CheckM [2] complete and contamination scores larger than or equal to 95.0 and smaller than or equal to 1.0, respectively.

### Supplemental Tables

**Supplemental Table S1.** Comparison of functional annotations and identification of coding sequences on 35 selected genomes from RefSeq

| Genome / Tool | Total CDS | Hypothetical CDS[a] | Identified CDS[b] | Total sORF[c] |
|---|---|---|---|---|
| *Pseudomonas aeruginosa* PAO1 (GCF_000006765.1) | | | | |
| Prokka | 5,671 | 38.3% (2,172) | - | - |
| DFAST | 5,727 | 33.5% (1,921) | - | - |
| Bakta | 5,685 | 5.8% (330) | 97.6 % (5,549) | 4 |
| Bakta w/o AFSI | 5,682 | 6.5% (372) | - | 1 |
| *Staphylococcus aureus subsp. aureus* NCTC 8325 (GCF_000013425.1) | | | | |
| Prokka | 2,630 | 20.6% (542) | - | - |
| DFAST | 2,676 | 17.5% (467) | - | - |
| Bakta | 2,644 | 4.8% (128) | 99.3% (2,626) | 13 |
| Bakta w/o AFSI | 2,636 | 6.2% (163) | - | 5 |
| *Staphylococcus epidermidis* ATCC 14990 (GCF_006094375.1) | | | | |
| Prokka | 2,259 | 20.6% (465) | - | - |
| DFAST | 2,294 | 19.7% (458) | - | - |
| Bakta | 2,264 | 7.4% (167) | 97.2% (2,200) | 5 |
| Bakta w/o AFSI | 2,263 | 8.1% (184) | - | 4 |
| *Enterococcus faecium* ATCC 8459 = NRRL B-2354 (GCF_000336405.1) | | | | |
| Prokka | 2,724 | 22.5% (613) | - | - |
| DFAST | 2,759 | 26.8% (740) | - | - |
| Bakta | 2,726 | 10.9% (298) | 96.7% (2,637) | 2 |
| Bakta w/o AFSI | 2,724 | 11.9% (325) | - | 0 |

| | | | | |
|---|---|---|---|---|
| *Klebsiella pneumoniae* ATCC BAA-2146 (GCF_000364385.3) | | | | |
| Prokka | 5,510 | 28.7% (1,584) | - | - |
| DFAST | 5,606 | 21.6% (1,211) | - | - |
| Bakta | 5,528 | 5.9% (328) | 98.6% (5,451) | 17 |
| Bakta w/o AFSI | 5,519 | 6.6% (363) | - | 10 |
| *Listeria monocytogenes* R2-502 (GCF_000438585.1) | | | | |
| Prokka | 3,028 | 38.6% (1,168) | - | - |
| DFAST | 3,054 | 24.8% (758) | - | - |
| Bakta | 3,036 | 7.2% (218) | 97.5% (2,959) | 5 |
| Bakta w/o AFSI | 3,035 | 8.0% (243) | - | 4 |
| *Escherichia coli* O157:H7 Sakai (GCF_000008865.2) | | | | |
| Prokka | 5,374 | 29.7% (1,598) | - | - |
| DFAST | 5,337 | 23.0% (1,229) | - | - |
| Bakta | 5,449 | 4.0% (216) | 99.8% (5,436) | 76 |
| Bakta w/o AFSI | 5,404 | 4.8% (261) | - | 32 |
| *Enterobacter cloacae* GGT036 (GCF_000770155.1) | | | | |
| Prokka | 4,479 | 24.3% (1,089) | - | - |
| DFAST | 4,496 | 15.5% (695) | - | - |
| Bakta | 4,491 | 4.3% (192) | 95.5% (4,287) | 11 |
| Bakta w/o AFSI | 4,488 | 4.7% (210) | - | 8 |
| *Salmonella enterica* subsp. *enterica* serovar Infantis 1326/28 (GCF_000953495.1) | | | | |
| Prokka | 4,360 | 23.7% (1,035) | - | - |
| DFAST | NA | NA (NA) | - | - |
| Bakta | 4,375 | 2.9% (126) | 97.0% (4,244) | 16 |
| Bakta w/o AFSI | 4,366 | 3.3% (146) | - | 8 |
| *Acinetobacter baumannii* ab736 (GCF_002116925.1) | | | | |
| Prokka | 3,712 | 43.3% (1,609) | - | - |
| DFAST | 3,711 | 28.0% (1,039) | - | - |

| | | | | |
|---|---|---|---|---|
| Bakta | 3,713 | 13.3% (493) | 99.9% (3,710) | 1 |
| Bakta w/o AFSI | 3,713 | 14.8% (548) | - | 1 |
| *Bacillus subtilis* subsp. *subtilis* 168 (GCF_000009045.1) | | | | |
| Prokka | 4,214 | 32.2% (1,358) | - | - |
| DFAST | 4,242 | 26.4% (1,119) | - | - |
| Bakta | 4,225 | 1.7% (72) | 97.8% (4,130) | 8 |
| Bakta w/o AFSI | 4,221 | 2.2% (91) | - | 6 |
| *Serratia marcescens* subsp. *marcescens* ATCC 13880 substr. Sm_S95_jyu2015 (GCF_017299435.1) | | | | |
| Prokka | 4,734 | 28.0% (1,324) | - | - |
| DFAST | 4,814 | 20.4% (983) | - | - |
| Bakta | 4,746 | 5.4% (257) | 96.7% (4,590) | 10 |
| Bakta w/o AFSI | 4,743 | 6.1% (291) | - | 7 |
| *Vibrio cholerae* MS6 (GCF_000829215.1) | | | | |
| Prokka | 3,550 | 35.7% (1,267) | - | - |
| DFAST | 3,549 | 17.6% (623) | - | - |
| Bakta | 3,556 | 5.8% (205) | 97.6% (3,469) | 4 |
| Bakta w/o AFSI | 3,553 | 6.3% (225) | - | 2 |
| *Vibrio parahaemolyticus* VPD14 (GCF_004006515.1) | | | | |
| Prokka | 4,561 | 40.1% (1,831) | - | - |
| DFAST | 4,549 | 19.0% (865) | - | - |
| Bakta | 4,567 | 9.9% (452) | 95.5% (4,362) | 4 |
| Bakta w/o AFSI | 4,567 | 10.3% (471) | - | 4 |
| *Borreliella burgdorferi* B31_NRZ (GCF_002151505.1) | | | | |
| Prokka | 1,339 | 63.1% (845) | - | - |
| DFAST | 1,352 | 44.4% (600) | - | - |
| Bakta | 1,339 | 12.4% (166) | 91.6% (1,226) | 0 |
| Bakta w/o AFSI | 1,339 | 14.1% (189) | - | 0 |

| | | | | |
|---|---|---|---|---|
| *Campylobacter jejuni* subsp. *jejuni* NCTC 11168 (GCF_000009085.1) | | | | |
| Prokka | 1,658 | 37.9% (629) | - | - |
| DFAST | 1,662 | 19.3% (321) | - | - |
| Bakta | 1,660 | 5.5% (91) | 95.1% (1,579) | 2 |
| Bakta w/o AFSI | 1,659 | 6.8% (112) | - | 1 |
| *Clostridioides difficile* BR81 (GCF_002007885.1) | | | | |
| Prokka | 3,592 | 41.7% (1,498) | - | - |
| DFAST | 3,649 | 20.3% (741) | - | - |
| Bakta | 3,596 | 6.1% (220) | 99.6% (3,581) | 4 |
| Bakta w/o AFSI | 3,593 | 7.3% (264) | - | 1 |
| *Clostridium botulinum* DFPST0029 (GCF_003058345.1) | | | | |
| Prokka | 3,475 | 43.1% (1,499) | - | - |
| DFAST | 3,521 | 24.6% (866) | - | - |
| Bakta | 3,478 | 8.9% (309) | 96.6% (3,359) | 2 |
| Bakta w/o AFSI | 3,476 | 10.2% (354) | - | 1 |
| *Clostridium tetani* 12124569 (GCF_000967115.2) | | | | |
| Prokka | 2,849 | 46.9% (1,335) | - | - |
| DFAST | 2,874 | 37.7% (1,084) | - | - |
| Bakta | 2,851 | 15.5% (441) | 84.2% (2,401) | 0 |
| Bakta w/o AFSI | 2,851 | 16.3% (464) | - | 0 |
| *Streptococcus pneumoniae* NT_110_58 (GCF_000817005.1) | | | | |
| Prokka | 2,246 | 45.3% (1,017) | - | - |
| DFAST | 2,254 | 31.3% (706) | - | - |
| Bakta | 2,267 | 7.1% (162) | 99.7% (2,258) | 20 |
| Bakta w/o AFSI | 2,265 | 9.6% (217) | - | 17 |
| *Streptococcus pyogenes* NGAS638 (GCF_001267845.1) | | | | |
| Prokka | 1,699 | 37.3% (634) | - | - |

| | | | | |
|---|---|---|---|---|
| DFAST | 1,687 | 34.3% (578) | - | - |
| Bakta | 1,703 | 5.3% (91) | 95.3% (1,624) | 4 |
| Bakta w/o AFSI | 1,701 | 6.5% (110) | - | 2 |
| *Corynebacterium diphtheriae* NCTC11397 (GCF_001457455.1) | | | | |
| Prokka | 2,328 | 45.4% (1,058) | - | - |
| DFAST | 2,316 | 37.4% (867) | - | - |
| Bakta | 2,328 | 14.5% (338) | 99.9% (2,326) | 0 |
| Bakta w/o AFSI | 2,328 | 15.0% (350) | - | 0 |
| *Treponema pallidum* subsp. *pallidum* X-4 (GCF_005885795.1) | | | | |
| Prokka | 978 | 45.9% (449) | - | - |
| DFAST | 982 | 38.7% (380) | - | - |
| Bakta | 978 | 6.2% (61) | 77.2% (755) | 0 |
| Bakta w/o AFSI | 978 | 7.2% (70) | - | 0 |
| *Helicobacter pylori* Puno135 (GCF_000224555.1) | | | | |
| Prokka | 1,537 | 45.1% (693) | - | - |
| DFAST | 1,538 | 28.0% (430) | - | - |
| Bakta | 1,541 | 11.2% (173) | 92.9% (1,431) | 4 |
| Bakta w/o AFSI | 1,540 | 12.0% (185) | - | 3 |
| *Rickettsia rickettsii* Iowa (GCF_000017445.4) | | | | |
| Prokka | 1,427 | 56.6% (808) | - | - |
| DFAST | 1,365 | 53.0% (723) | - | - |
| Bakta | 1,428 | 28.4% (406) | 87.5% (1,249) | 0 |
| Bakta w/o AFSI | 1,428 | 29.6% (422) | - | 0 |
| *Legionella pneumophila* L10-023 (GCF_001610735.1) | | | | |
| Prokka | 3,533 | 52.3% (1,848) | - | - |
| DFAST | 3,471 | 36.3% (1,261) | - | - |
| Bakta | 3,352 | 17.8% (628) | 92.0% (3,085) | 0 |
| Bakta w/o AFSI | 3,532 | 18.7% (660) | - | 0 |

| | | | | |
|---|---|---|---|---|
| *Yersinia enterocolitica* FORC_002 (GCF_000987925.1) | | | | |
| Prokka | 4,327 | 30.3% (1,313) | - | - |
| DFAST | 4,306 | 20.2% (868) | - | - |
| Bakta | 4,338 | 7.6% (328) | 93.8% (4,068) | 8 |
| Bakta w/o AFSI | 4,336 | 7.8% (338) | - | 6 |
| *Bacillus thuringiensis* ATCC 10792 (GCF_002119445.1) | | | | |
| Prokka | 6,327 | 50.8% (3,211) | - | - |
| DFAST | 6,388 | 36.2% (2,315) | - | - |
| Bakta | 6,341 | 12.7% (807) | 98.9% (6,271) | 8 |
| Bakta w/o AFSI | 6,339 | 14.7% (931) | - | 5 |
| *Lactobacillus casei* DSM 20011 (GCF_000829055.1) | | | | |
| Prokka | 2,890 | 44.6% (1,288) | - | - |
| DFAST | 2,905 | 34.1% (990) | - | - |
| Bakta | 2,893 | 13.7% (397) | 92.7% (2,681) | 0 |
| Bakta w/o AFSI | 2,891 | 14.7% (425) | - | 0 |
| *Streptomyces antibioticus* DSM 41481 (GCF_012044595.1) | | | | |
| Prokka | 7,612 | 55.6% (4,231) | - | - |
| DFAST | 7,831 | 41.3% (3,233) | - | - |
| Bakta | 7,620 | 16.7% (1,276) | 82.5% (6,287) | 1 |
| Bakta w/o AFSI | 7,619 | 17.5% (1,335) | - | 0 |
| *Corynebacterium jeikeium* NCTC11914 (GCF_900477975.1) | | | | |
| Prokka | 2,057 | 40.7% (838) | - | - |
| DFAST | 2,078 | 34.0% (707) | - | - |
| Bakta | 2,060 | 8.3% (170) | 99.8% (2,055) | 2 |
| Bakta w/o AFSI | 2,059 | 10.2% (209) | - | 1 |
| *Xanthomonas campestris* M28 (GCF_014841015.1) | | | | |
| Prokka | 4,326 | 46.7% (2,022) | - | - |
| DFAST | 4,358 | 29.2% (1,274) | - | - |

| | | | | |
|---|---|---|---|---|
| Bakta | 4,333 | 12.9% (559) | 84.6% (3,664) | 2 |
| Bakta w/o AFSI | 4,332 | 13.9% (600) | - | 1 |
| *Sinorhizobium meliloti* SM11 (GCF_000218265.1) | | | | |
| Prokka | 6,913 | 47.7% (3,298) | - | - |
| DFAST | 6,927 | 29.2% (2,025) | - | - |
| Bakta | 6,920 | 14.4% (998) | 94.6% (6,547) | 2 |
| Bakta w/o AFSI | 6,920 | 15.3% (1,058) | - | 2 |
| *Sorangium cellulosum* So ce 56 (GCF_000067165.1) | | | | |
| Prokka | 9,627 | 65.4% (6,297) | - | - |
| DFAST | 9,933 | 70.6% (7,012) | - | - |
| Bakta | 9,654 | 25.6% (2,473) | 86.2% (8,326) | 0 |
| Bakta w/o AFSI | 9,654 | 26.4% (2,549) | - | 0 |
| *Thiobacillus denitrificans* ATCC 25259 (GCF_000012745.1) | | | | |
| Prokka | 2,814 | 42.3% (1,190) | - | - |
| DFAST | 2,842 | 36.6% (1,039) | - | - |
| Bakta | 2,818 | 11.6% (326) | 95.2% (2,682) | 0 |
| Bakta w/o AFSI | 2,818 | 11.7% (330) | - | 0 |
| **Total** | | | | |
| Prokka | 130,360 | 41.2% (53,656) | - | - |
| DFAST | 127,053 | 31.6% (40,128) | - | - |
| Bakta | 130,683 | 10.6% (13,902) | 94.2% (123,105) | 235 |
| Bakta w/o AFSI | 130,573 | 11.5% (15,066) | - | 132 |

Note: Prokka, DFAST and Bakta were executed with default parameters providing the following information relevant to the annotation workflow: genus, species and assembly status. A detailed list of command lines is provided in Supplemental Notes S3; Bakta w/o AFSI: Bakta without alignment-free sequence identification

[a] Protein sequences of unknown function, *i.e.* product denoted as either *hypothetical protein*, *putative protein, uncharacterized protein* or *conserved predicted protein*.

[b] CDS identified with assigned public database identifiers from either RefSeq, UniParc or UniRef.

[c] Short ORFs shorter than 29 amino acids

**Supplemental Table S2.** Functional annotation and protein identification benchmark set comprising 362 selected genomes from GenBank

| GenBank accessions |
|---|
| GCA_018141845.1, GCA_016865485.1, GCA_016865505.1, GCA_018228645.1, GCA_018228685.1, |
| GCA_019203805.1, GCA_902502875.2, GCA_014191545.1, GCA_018127705.1, GCA_009497875.2, |
| GCA_013467635.1, GCA_013467655.1, GCA_015476115.1, GCA_017948305.1, GCA_018672315.1, |
| GCA_017798125.1, GCA_018866245.1, GCA_015243575.1, GCA_016458825.1, GCA_017569285.1, |
| GCA_018118585.1, GCA_018128945.1, GCA_016865015.1, GCA_016865035.1, GCA_017134815.1, |
| GCA_015693905.1, GCA_016887485.1, GCA_016350165.1, GCA_016887505.1, GCA_003352025.1, |
| GCA_018394295.1, GCA_018394315.1, GCA_018622495.1, GCA_018622675.1, GCA_018069705.1, |
| GCA_018398915.1, GCA_004376055.2, GCA_017798145.1, GCA_017309585.1, GCA_015482585.1, |
| GCA_014218215.1, GCA_013457415.1, GCA_017161075.1, GCA_017945865.1, GCA_017751145.1, |
| GCA_016806125.1, GCA_010681825.2, GCA_019355395.1, GCA_014023125.1, GCA_015099395.1, |
| GCA_014873095.1, GCA_013745115.1, GCA_016865445.1, GCA_017357865.1, GCA_013874835.1, |
| GCA_014041935.1, GCA_014337215.1, GCA_014337195.1, GCA_014337155.1, GCA_014337175.1, |
| GCA_016126655.1, GCA_016591975.1, GCA_015679245.1, GCA_017808555.1, GCA_017569265.1, |
| GCA_017569305.1, GCA_016894345.1, GCA_016925615.1, GCA_016916535.1, GCA_017498545.1, |
| GCA_017569225.1, GCA_018437505.1, GCA_017945845.1, GCA_907165195.1, GCA_019285715.1, |
| GCA_018604165.1, GCA_019195455.1, GCA_018916745.1, GCA_014654535.1, GCA_000177275.2, |
| GCA_000463875.1, GCA_000419805.1, GCA_010206225.2, GCA_000494665.1, GCA_000494645.1, |
| GCA_001657615.1, GCA_001644275.1, GCA_001644325.1, GCA_001644345.1, GCA_001644365.1, |
| GCA_001644415.1, GCA_001644495.1, GCA_001644255.1, GCA_001644445.1, GCA_001644455.1, |
| GCA_001644245.1, GCA_015207155.1, GCA_015207075.1, GCA_015207195.1, GCA_015206905.1, |
| GCA_015207365.1, GCA_014132315.1, GCA_003003915.1, GCA_015233825.1, GCA_009915105.1, |
| GCA_018326565.1, GCA_018326585.1, GCA_018326665.1, GCA_018326685.1, GCA_018326705.1, |
| GCA_018326725.1, GCA_018326745.1, GCA_018326765.1, GCA_018326925.1, GCA_018327005.1, |
| GCA_018327065.1, GCA_018327085.1, GCA_018327145.1, GCA_018327205.1, GCA_018327385.1, |
| GCA_018327405.1, GCA_018327445.1, GCA_018327465.1, GCA_015223225.1, GCA_015234195.1, |
| GCA_009767105.1, GCA_015700665.1, GCA_015700695.1, GCA_010692825.1, GCA_014695345.1, |
| GCA_014695275.1, GCA_014695325.1, GCA_014695255.1, GCA_014695195.1, GCA_009900665.1, |
| GCA_009908265.2, GCA_018449595.1, GCA_018449715.1, GCA_014892285.1, GCA_014892275.1, |
| GCA_014892335.1, GCA_014892345.1, GCA_014892395.1, GCA_014892495.1, GCA_904424815.1, |
| GCA_015863215.1, GCA_015863195.1, GCA_015863185.1, GCA_014174405.1, GCA_014116815.1, |
| GCA_016123485.1, GCA_014384745.1, GCA_014384795.1, GCA_014384805.1, GCA_014384785.1, |
| GCA_014384885.1, GCA_014384905.1, GCA_014384765.1, GCA_014384895.1, GCA_014384705.1, |
| GCA_014385265.1, GCA_014384965.1, GCA_014384925.2, GCA_014384995.1, GCA_014385225.1, |
| GCA_014384985.1, GCA_014385005.1, GCA_014385025.1, GCA_014385135.2, GCA_014385165.1, |

GCA_014385105.1, GCA_014385095.1, GCA_014385085.1, GCA_014385195.1, GCA_014306095.1, GCA_015472025.1, GCA_014779695.1, GCA_014858005.1, GCA_014841525.1, GCA_014841595.1, GCA_014841765.1, GCA_014841835.1, GCA_014841995.1, GCA_014842315.1, GCA_014857905.1, GCA_015206945.1, GCA_015206935.1, GCA_015207035.1, GCA_016587355.1, GCA_016587455.1, GCA_014982975.1, GCA_014982785.1, GCA_014982745.1, GCA_014982715.1, GCA_014983025.1, GCA_015476615.1, GCA_015546655.1, GCA_018524345.1, GCA_015689585.1, GCA_015689535.1, GCA_015689545.1, GCA_015689575.1, GCA_015689655.1, GCA_015689635.1, GCA_015689685.1, GCA_015689645.1, GCA_015689705.1, GCA_015689745.1, GCA_015689735.1, GCA_015689785.1, GCA_015689775.1, GCA_015689805.1, GCA_016464625.1, GCA_016522115.1, GCA_016458225.1, GCA_016595455.1, GCA_016595575.1, GCA_016811075.1, GCA_016892645.1, GCA_016892445.1, GCA_016918765.1, GCA_018332375.1, GCA_017948655.1, GCA_018348555.1, GCA_018348615.1, GCA_018348635.1, GCA_018682635.1, GCA_017114885.1, GCA_017942165.1, GCA_018145715.1, GCA_018122655.1, GCA_018223375.1, GCA_018223435.1, GCA_018362825.1, GCA_018362815.1, GCA_018362855.1, GCA_018413565.1, GCA_019083985.1, GCA_018982925.1, GCA_019008365.1, GCA_018883545.1, GCA_018919725.1, GCA_018919645.1, GCA_018919585.1, GCA_018919625.1, GCA_018919565.1, GCA_019084065.1, GCA_019061145.1, GCA_019084025.1, GCA_019132875.1, GCA_019166125.1, GCA_019218305.1, GCA_019218315.1, GCA_019218365.1, GCA_019218375.1, GCA_019218415.1, GCA_019218445.1, GCA_015550775.1, GCA_015551295.1, GCA_015553565.1, GCA_015554585.1, GCA_015557515.1, GCA_015559035.1, GCA_015560015.1, GCA_015669665.1, GCA_018784685.1, GCA_018784725.1, GCA_015555245.1, GCA_015559165.1, GCA_015559785.1, GCA_015561015.1, GCA_015667115.1, GCA_000428245.1, GCA_011800315.1, GCA_015207585.1, GCA_013519935.1, GCA_003426325.1, GCA_003426525.1, GCA_003426555.1, GCA_003585675.1, GCA_002968105.1, GCA_002968095.1, GCA_003030805.1, GCA_014610835.1, GCA_014156375.1, GCA_004791555.1, GCA_018326505.1, GCA_018326525.1, GCA_018326545.1, GCA_018326625.1, GCA_018326645.1, GCA_018326785.1, GCA_018326805.1, GCA_018326825.1, GCA_018326845.1, GCA_018326865.1, GCA_018326885.1, GCA_018326905.1, GCA_018326945.1, GCA_018326965.1, GCA_018326985.1, GCA_018327025.1, GCA_018327045.1, GCA_018327105.1, GCA_018327125.1, GCA_018327165.1, GCA_018327185.1, GCA_018327225.1, GCA_018327245.1, GCA_018327265.1, GCA_018327285.1, GCA_018327305.1, GCA_018327325.1, GCA_018327345.1, GCA_018327365.1, GCA_018327425.1, GCA_018784815.1, GCA_014873415.1, GCA_014873965.1, GCA_016467405.1, GCA_015700715.1, GCA_016617495.1, GCA_016617515.1, GCA_014529625.1, GCA_014529675.1, GCA_014529665.1, GCA_015354435.1, GCA_015354475.1, GCA_017898095.1, GCA_014837105.1, GCA_014837035.1, GCA_014836765.1, GCA_015666175.1, GCA_018598225.1, GCA_018598175.1, GCA_018598585.1, GCA_018598185.1, GCA_015905305.1, GCA_015905225.1, GCA_015905245.1, GCA_015550005.1, GCA_015669005.1, GCA_016649355.1, GCA_016649365.1, GCA_016649425.1, GCA_017810195.1, GCA_016521575.1, GCA_016469015.1, GCA_016632365.1, GCA_016918935.1, GCA_019140855.1, GCA_019148575.1, GCA_017948555.1, GCA_018128545.1, GCA_017592665.1, GCA_018223385.1, GCA_018223415.1, GCA_018223365.1, GCA_018704065.1, GCA_018729455.1, GCA_019042235.1, GCA_019042245.1, GCA_019042275.1

**Supplemental Table S3.** Functional annotation and protein identification benchmark set comprising 198 selected metagenome-assembled genomes from GenBanks

| GenBank accession |
|---|
| GCA_002297765.1, GCA_002298295.1, GCA_002298975.1, GCA_002298225.1, GCA_002297475.1, GCA_002295925.1, GCA_002293145.1, GCA_002314135.1, GCA_002311805.1, GCA_002308695.1, GCA_002307725.1, GCA_002306875.1, GCA_002306465.1, GCA_002306155.1, GCA_002306335.1, GCA_002305805.1, GCA_002325565.1, GCA_002325245.1, GCA_002325045.1, GCA_002322275.1, GCA_002321895.1, GCA_002320175.1, GCA_002320085.1, GCA_002320055.1, GCA_002320005.1, GCA_002337405.1, GCA_002337285.1, GCA_002337145.1, GCA_002337125.1, GCA_002336245.1, GCA_002335085.1, GCA_002335425.1, GCA_002333005.1, GCA_002332985.1, GCA_002331575.1, GCA_002331385.1, GCA_002329025.1, GCA_002328995.1, GCA_002328335.1, GCA_002328165.1, GCA_002347645.1, GCA_002345215.1, GCA_002345185.1, GCA_002345145.1, GCA_002344965.1, GCA_002344785.1, GCA_002344345.1, GCA_002343985.1, GCA_002342295.1, GCA_002341725.1, GCA_002341585.1, GCA_002341545.1, GCA_002341485.1, GCA_002340805.1, GCA_002340585.1, GCA_002340255.1, GCA_002359515.1, GCA_002359395.1, GCA_002359265.1, GCA_002359215.1, GCA_002359185.1, GCA_002354635.1, GCA_002354605.1, GCA_002353655.1, GCA_002352555.1, GCA_002352105.1, GCA_002348315.1, GCA_002346225.1, GCA_002367555.1, GCA_002366545.1, GCA_002365175.1, GCA_002364745.1, GCA_002364215.1, GCA_002364085.1, GCA_002366485.1, GCA_002378905.1, GCA_002378005.1, GCA_002376875.1, GCA_002376725.1, GCA_002376625.1, GCA_002376045.1, GCA_002375365.1, GCA_002375495.1, GCA_002375465.1, GCA_002373505.1, GCA_002373475.1, GCA_002371595.1, GCA_002372835.1, GCA_002369135.1, GCA_002393145.1, GCA_002392595.1, GCA_002392045.1, GCA_002391865.1, GCA_002385275.1, GCA_002385225.1, GCA_002385025.1, GCA_002384665.1, GCA_002384935.1, GCA_002382325.1, GCA_002382705.1, GCA_002381615.1, GCA_002380645.1, GCA_002381125.1, GCA_002395615.1, GCA_002394385.1, GCA_002389905.1, GCA_002390845.1, GCA_002400415.1, GCA_002389085.1, GCA_002389665.1, GCA_002388685.1, GCA_002388665.1, GCA_002387375.1, GCA_002386415.1, GCA_002406015.1, GCA_002405475.1, GCA_002405165.1, GCA_002403705.1, GCA_002403465.1, GCA_002403435.1, GCA_002403335.1, GCA_002402585.1, GCA_002399785.1, GCA_002399025.1, GCA_002397905.1, GCA_002415215.1, GCA_002414885.1, GCA_002415705.1, GCA_002415665.1, GCA_002414525.1, GCA_002415565.1, GCA_002414445.1, GCA_002414385.1, GCA_002415385.1, GCA_002413005.1, GCA_002412965.1, GCA_002410345.1, GCA_002410145.1, GCA_002424395.1, GCA_002427365.1, GCA_002424235.1, GCA_002426725.1, GCA_002425805.1, GCA_002425185.1, GCA_002421065.1, GCA_002420425.1, GCA_002420525.1, GCA_002420305.1, GCA_002419775.1, GCA_002419535.1, GCA_002418285.1, GCA_002418245.1, GCA_002418095.1, GCA_002432845.1, GCA_002431665.1, GCA_002431625.1, GCA_002430265.1, GCA_002427895.1, GCA_002427815.1, GCA_002427755.1, GCA_002428995.1, GCA_002435785.1, GCA_002440635.1, GCA_002439045.1, GCA_002438025.1, GCA_002437725.1, GCA_002439815.1, GCA_002436875.1, GCA_002433365.1, GCA_002456035.1, |

| |
|---|
| GCA_002455715.1, GCA_002454815.1, GCA_002453945.1, GCA_002452095.1, GCA_002449395.1, GCA_002450875.1, GCA_002449845.1, GCA_002492145.1, GCA_002471845.1, GCA_002471675.1, GCA_002471575.1, GCA_002471335.1, GCA_002471125.1, GCA_002470515.1, GCA_002476985.1, GCA_002479255.1, GCA_002477825.1, GCA_002484125.1, GCA_002484005.1, GCA_002483535.1, GCA_002482645.1, GCA_002482605.1, GCA_002482265.1, GCA_002480685.1, GCA_002480575.1, GCA_002500565.1, GCA_002501065.1, GCA_002499995.1 |

### Supplemental Notes

**Supplemental Notes S1**

During the SQLite database initialization procedure, UPSs, IPSs and PSCs records are created, importing unique protein sequences from UniRef100 and UniParc and protein cluster seed sequences from UniRef90 [3]. Subsequently, these database records are refined with annotations from additional external databases. Protein products and gene symbols are extracted from RefSeq non-redundant protein records [4], UniProt/SwissProt [3], AMRFinderPlus [5] or ISfinder [6]. Furthermore, annotations are enriched with additional information like EC numbers, COG functional categories [7] and GO [8] terms. All annotations are conducted and supersede each other in the outlined order according to the specificity of annotation sources.

1. RefSeq non-redundant proteins: UPSs are annotated with identifiers of RefSeq non-redundant protein records which are merged via PSHDs. IPSs or related PSCs are annotated with gene symbols.
2. NCBI COG: PSCs are annotated with COG function and category records via a large-scale homology search against cluster representative sequences of the COG [7] database using Diamond [9] applying a mutual coverage threshold 0.80 and a sequence identity threshold of 0.90. PSCs without a gene symbol or protein product different from *hypothetical protein* are annotated accordingly.
3. UniProt/SwissProt: IPSs and PSCs are annotated with gene symbols, protein products, EC numbers and GO terms from related SwissProt [3] records identified via PSHDs.
4. NCBI AMRFinderPlus: IPSs are annotated with gene symbols and protein products via PSHDs. PSCs are annotated with gene symbols and protein products by HMMER [10] hits against AMRFinderPlus HMMs.
5. ISfinder: IPSs and PSCs are annotated with gene symbols and protein products via large-scale homology searches against transposase protein sequences using Diamond [9] with a mutual coverage threshold of 0.99 and 0.80 and a sequence identity threshold of 0.98 and 0.90 for IPSs and PSCs, respectively.

**Supplemental Notes S2**

To provide high-quality annotation for certain genes of special interest that might not be able to sufficiently annotate via protein sequence hash digests or protein cluster sequences, Bakta takes advantage of a protein sequence-based expert annotation system integrating high-quality annotation sources from external databases. Therefore, protein sequences, gene symbols, protein products, query and subject coverage thresholds, sequence identity thresholds and priority ranks are stored for protein sequences from VFDB [11] and NCBI BlastRules [4]. To give precedence on more precise annotations in case of multiple annotations, these different sources are given a rank taking into account proprietary priority levels:

NCBI BlastRules BlastRuleEquivalog: 70
VFDB core sequences: 75
NCBI BlastRules BlastRuleException: 80
NCBI BlastRules BlastRuleIS: 90
NCBI BlastRules BlastRuleExact: 99

**Supplemental Notes S3**

Prokka, DFAST and Bakta were executed with the following command line options:

Prokka

```
prokka --outdir . --force --prefix <accession> --genus <genus>
--species <species> --strain <strain> --gcode 11 --usegenus --cpus 8
--rfam <input>
```

DFAST

```
dfast --out output --organism "<genus> <species>" --strain <strain>
--use_original_name t --sort_sequence f --cpu 8 --genome <input>
```

Bakta

```
bakta --db <baktadb> --prefix <prefix> --genus <genus> --species
<species> --strain <strain> --keep-contig-headers --threads 8
--complete <input>
```

**Supplemental Notes S4**

The backend of our Bakta web application is based on a scalable REST-API implemented in Go. All input data is isolated on a per-job level, by the provision of dedicated cloud based object storage endpoints, *i.e.* buckets, for each annotation job. Actual annotations are conducted by processes of the Bakta command line version distributed on a scalable Kubernetes cluster. Currently, this Kubernetes cluster is hosted and run within a cloud computing infrastructure of the de.NBI consortium. Hence, required hardware resources can be seamlessly scaled to current demands. Besides accepting jobs from our web application, this public API can also be used independently as an annotation microservice, *e.g.* for the integration into larger web-based analysis pipelines. A comprehensive documentation of this API is provided at https://bakta.readthedocs.io.

8;47(D1):D687–92. Available from: http://dx.doi.org/10.1093/nar/gky1080
